# Oviposition behavior, nymphal development, and below-ground olfactory preference of *Pentastiridius leporinus* on twelve *Beta vulgaris* genotypes

**DOI:** 10.1101/2025.10.02.679987

**Authors:** Pamela Bruno, Shahinoor Rahman, Karthi Balakrishnan, Britta Kais, Jürgen Gross, Michael Rostás

**Author notes:** Corresponding authors: Pamela Bruno, Michael Rostás. Authors contributed equally (Pamela Bruno, Shahinoor Rahman). email addresses: Shahinoor Rahman, Karthi Balakrishnan, Britta Kais, Jürgen Gross.

## Abstract

The planthopper *Pentastiridius leporinus* (Hemiptera: Cixiidae) is the primary vector of two phloem-restricted prokaryotes, which can cause the disease syndrome de basses richesses (SBR) in sugar beet (*Beta vulgaris* L.). Heavy infestations of this insect have contributed to significant yield losses in Central Europe, but little is known about how different *B. vulgaris* genotypes influence vector behavior and performance. Understanding such interactions is critical for both pest management and the development of sugar beet cultivars that are less favorable to the vector. We investigated oviposition decisions, nymphal development, and belowground chemotaxis of *P. leporinus* using twelve *B. vulgaris* genotypes: eight sugar beet genotypes with contrasting SBR tolerance, two Swiss chard, and two beetroot varieties. In no-choice assays, females oviposited at similar rates on all tested genotypes, indicating broad host suitability. However, in three-choice tests, significant differences emerged, with several genotypes receiving more egg batches than the reference variety Vasco. Despite adult host preferences, nymphal survival, growth, and instar development did not differ among genotypes. In contrast, belowground olfactometer assays revealed that nymphs exhibited clear chemotactic responses: several genotypes attracted significantly more individuals than others. This represents the first documentation of chemotactic host location in *P. leporinus* nymphs. Our findings reveal a mismatch between adult oviposition preference and nymphal performance, a pattern not uncommon in other generalist herbivores. Both adult and nymphal behaviors were nevertheless influenced by genotype-specific traits, likely linked to plant chemistry. Identifying these cues could inform breeding strategies aimed at reducing vector attraction, thereby complementing pathogen-targeted approaches and supporting integrated management of SBR in sugar beet.

## Introduction

Sugar beet (*Beta vulgaris* L.) is a major crop for sugar production worldwide, but it faces serious threats from syndrome de basses richesses (SBR), a disease mainly caused by the phloem-restricted γ-3 proteobacterium *Candidatus* Arsenophonus phytopathogenicus and the phytoplasma *Candidatus* Phytoplasma solani (Gatineau et al., 2002; Sémétey, Gatineau, et al., 2007; Lang et al. 2025). SBR stunts plant growth, induces nutrient deficiencies, and leads to significant yield losses, posing a major economic challenge for the sugar beet industry (Agyei et al., 2025; Pfitzer et al., 2020). The spread of SBR has been particularly noted in Germany and Switzerland, where rapid spread of *P. leporinus* and the lack of effective control measures for both the pathogen and its vector have led to substantial economic losses (Behrmann et al., 2021; Lang, 2025; Pfitzer et al., 2024; Pfitzer et al., 2020).

The primary vector of the pathogen in sugar beet is *Pentastiridius leporinus* (Hemiptera: Cixiidae), a planthopper whose ecological interactions with the host plant and vector competence are beginning to be elucidated (Behrmann et al., 2022; Bressan et al., 2009; Bressan et al., 2008; Pfitzer et al., 2022; Sémétey, Bressan, et al., 2007). The planthopper’s ability to transmit the pathogen to other crops, such as potatoes, carrots and maybe onions, further complicates management strategies (Behrmann et al., 2023; Rinklef et al., 2024; Therhaag, Schneider, et al., 2024; Therhaag, Ulrich, et al., 2024; Witczak et al., 2025). Understanding the biology and life cycle of *P. leporinus* is crucial for developing effective pest management strategies to mitigate the impact of SBR on sugar beet production.

Efforts to combat SBR in sugar beet have focused on breeding tolerant varieties that mitigate the impact of the pathogen (Laufer et al., 2024). Such strategies have primarily aimed at reducing the severity of disease symptoms and minimizing yield losses, with several commercial breeding programs achieving progress in developing SBR-tolerant genotypes (KWS, 2023; Pfitzer et al., 2020; SESVanderHave, 2021). However, to date, no breeding initiatives have directly targeted resistance to the vector, *P. leporinus*. Developing sugar beet varieties that deter vector feeding, disrupt nymphal development, or otherwise reduce vector preference could complement existing pathogen-targeted approaches. Integrating resistance against both the pathogen and its vector would provide a more comprehensive strategy to manage SBR.

This study examines the interactions between *P. leporinus* and different *B. vulgaris* genotypes, namely eight genotypes of sugar beet (*B. vulgaris* subsp. *vulgaris*, cultivar group Altissima), two genotypes of Swiss chard (*B. vulgaris* subsp. *vulgaris*, cultivar group Flavescens) and two genotypes of beetroot (*B. vulgaris* subsp. *vulgaris*, cultivar group Conditiva). We focus on oviposition behavior of adult insects, nymphal growth, and their below-ground olfactory preference. By comparing these interactions, we aim to improve the understanding of the chemical ecology of *P. leporinus* as a vector of SBR, and contribute to the development of more tolerant sugar beet cultivars.

## Materials and Methods

### Insects

A rearing of *Pentastiridius leporinus* was established in the laboratory of Agricultural Entomology at the Georg-August University in Göttingen, DE, as detailed by Pfitzer et al. (2022). Briefly, adult *P. leporinus* were kept in 60 × 60 × 60 cm tents (BugDorm®, MegaView Science Co. Ltd, TW) containing one-month-old potted sugar beet plants, where they fed, reproduced, and oviposited. Egg batches were collected weekly and transferred to plastic boxes filled with a 3:1 mixture of peat (Fruhstorfer Erde Typ P 25, Hawita, DE) and sand (0–2 mm ø), together with a piece of sugar beet taproot. Nymphs fed on the root pieces, which were replaced weekly. Fifth-instar nymphs were moved to new pots with sugar beet plants and maintained there until adult emergence, after which they were returned to the oviposition tents. Newly-emerged adults (4 days old) were used for no-choice and choice oviposition experiments. Second-instar nymphs were used for nymph-performance experiments. Third-instar nymphs were used for belowground olfactometer assays. The nymphal stage was identified by measuring head capsule width, while the infection status within the rearing population was tested using nested polymerase chain reaction (Pfitzer et al., 2022). The infection rate was confirmed to be approximately 80% for ‘*Ca*. Arsenophonus phytopathogenicus’ whereas no infection was detected for ‘*Ca*. Phytoplasma solani’.

### Plant Material

Sugar beet of the variety “Vasco” (SESVanderHave Deutschland GmbH, Germany) was used for rearing *P. leporinus*. Additionally, four breeding companies each supplied two sugar beet genotypes, one presumed tolerant and one presumed susceptible, based on their expected response to SBR. The genotypes were labeled as follows: KWS SAAT SE & Co. KGaA (S1-S, S1-T), SESVanderHave (S2-S, S2-T), Hilleshög AB (S3-S, S3-T), and Strube D&S GmbH (S4-S, S4-T), where “S” indicates susceptibility and “T” indicates tolerance. Furthermore, four commercial varieties of Swiss chard and beetroot, obtained from KWS, were included in the experiments: the red Swiss chard “Fluence” (C1-r), the green Swiss Chard “Prius” (C2-g), and red beetroot varieties “Scarlett” (B1) and “Jolie” (B2). The inclusion of both red and green Swiss chard varieties allowed for potential color preferences.

Seeds were germinated in pots with a 3:1 mixture of peat (Fruhstorfer Erde Typ P 25, Hawita, Germany) and sand (0-2mm ø), and 2-week-old seedlings were transferred individually to pots with the same soil mixture. For performance experiments, 700 mL disposable polyethylene drink cups were used (Benail, United States), with perforations on the bottom and filled with 2 cm of sand and approx. 10 cm of soil mixture. A 5 cm layer of expanded clay was placed on top to provide a porous habitat for the nymphs. Plants used in the belowground olfactometer assays were grown in 170 mL polystyrene cylinders (Zuchtgläschen, K-TK e.K., Retzstadt, Germany). For the oviposition experiments, plants were grown in 500 mL transparent cylindrical pots made of polypropylene (PP), allowing egg batches to be monitored without disturbing the plants. Two-month-old plants were used for oviposition experiments, and six-week-old plants were used for performance and olfactometer experiments. All plants were grown in greenhouse conditions (18 – 24 °C, 65% RH, light: dark 16:8), watered and fertilized weekly with 1 g/L of ‘Hakaphos® Blau 15-10-15(+2)’ (COMPO EXPERT GmbH, Münster, Germany).

### Oviposition assays

Oviposition preferences of *P. leporinus* were tested under no-choice and three-choice conditions using 60 × 60 × 60 cm tents (BugDorm®, MegaView Science Co. Ltd, TW) with potted plants of different genotypes. In the no-choice test, each tent contained a single genotype, while in the three-choice test, three pots with different genotypes were placed together in one tent. In this case, sugar beet cultivar Vasco served as the baseline, since *P. leporinus* had been reared on this cultivar. In addition to Vasco, each tent was supplied with a pair of genotypes from one of the four breeding companies or a pair of Swiss chard or beetroot varieties. For both tests, groups of 10 females and 5 males, all four days old, were released into the tents and kept under controlled conditions (22 ± 2 °C, 65% RH). After 18 days, the number of egg batches per plant was recorded. For graphical representation (Fig. 2), oviposition was standardized using the sugar beet variety Vasco (the genotype routinely used for insect rearing) as the reference (set to 100%). Oviposition on all other genotypes was quantified as a relative oviposition index, calculated for each replicate as follows:

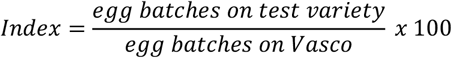

### Development of *P. leporinus* nymphs on *B. vulgaris* genotypes

To assess the performance of *P. leporinus* nymphs on different *B. vulgaris* genotypes, second-instar nymphs were individually placed on plants representing each of the 12 genotypes. Ten nymphs were carefully transferred to a plastic cup containing a single plant, and the cups were randomly arranged on plastic trays. All setups were maintained in a rearing chamber at 22 °C under a 16 : 8 h light : dark photoperiod. Plants were watered and fertilized weekly. To prevent the escape of emerging adults, each plant canopy was enclosed within a perforated clear polypropylene bag secured to the pot with a rubber band. The experiment was terminated upon the emergence of the first adult (90 days). At this time, surviving nymphs were collected, counted, and weighed collectively. Mean nymph weight and mortality rate per pot were calculated for each genotype. Each experimental setup was replicated 12 times.

### Belowground olfactometer tests

A four-arm belowground olfactometer (Rasmann et al., 2005; Rostás et al., 2015) was used to assess nymphal olfactory preferences. The central glass chamber (8 cm ø, 11 cm height) connected to four smaller chambers (5 cm ø, 11 cm height) via substrate-filled passages.

Plants grown in 170 mL pots were transferred with soil into the smaller chambers. The four test chambers of the olfactometer contained each a i) sugar beet plant of Vasco, ii) an SBR susceptible genotype, iii) an SBR tolerant genotype, iv) soil without plant. The contrasting sugar beet genotypes belonged to the same breeding company. In addition, both cultivars of beetroot and Swiss chard were tested in the same manner with Vasco as positive and soil as negative controls. The central chamber was filled with expanded clay granulate (2–5 mm, Pflanzgranulat, ASB Grünland Helmut Aurenz GmbH) up to a height of 6 cm, while the connecting passages were completely filled. Ten 3^rd^ instar nymphs were randomly selected from rearing boxes and then introduced into the central chamber. To prevent light entering the olfactometer arms, setups were covered with black cloth. After 24 hours, their positions were recorded; nymphs crossing into odor-containing arms were considered responsive. Positions of odor sources were randomized daily, and five olfactometers were run concurrently. Following each assay, glass components were washed with water, cleaned with acetone, dried under a fume hood, heated to 110°C for 1 hour, and cooled at room temperature. Room conditions were maintained at 22 ± 2°C and 50% relative humidity. For the graphical representation (Fig. 4), the genotype Vasco was used as the baseline reference and set to 100% based on the number of nymphs responding to it in each olfactometer assay. The values for the other two genotypes and the empty arm were then calculated relative to Vasco, to express their differences as increases or decreases in index values. A total of 20 replicates were conducted.

### Statistical analyses

Statistical analyses were performed using R software (v. 4.2.3; R Development Core Team, 2023). Analysis of variance (ANOVA) was conducted, followed by residual analysis to assess the fit of the models. Oviposition data across all *B. vulgaris* genotypes and from the 3-choice oviposition test (number of egg batches laid per treatment) were analyzed using generalized linear models (GLM) with a Poisson distribution. Nymphal weight differences were analyzed with generalized linear models (GLM) assuming a Gaussian distribution. The comparisons of instar achieved by the nymphs in the performance experiment was analyzed with GLM under Poisson distributions, and nymphal mortality across different *B. vulgaris* genotypes was compared using GLM with a binomial distribution. All analyses were followed by pairwise comparisons of estimated marginal means (emmeans) with Tukey adjustment for multiple testing. To analyze belowground olfactometer test results, nymph responses to the four different odor sources were analyzed using a general linear model (GLM) with binomial family distribution, considering number of successes (= all nymphs that went to each odor source) and number of failures (= all nymphs that went to the other three odor sources) as two-vector response variables. To account for the grouping of nymph choices released simultaneously with the same set of plants, a generalized linear mixed model (GLMM) including experimental day as a random effect was initially fitted. As the random effect did not significantly improve model fit, the simpler GLM without random effects was retained for analysis.

## Results

### Oviposition no-choice test

In the no-choice test, *P. leporinus* females laid eggs on all tested *B. vulgaris* genotypes at similar rates (Fig 1). No significant differences were observed in the number of egg batches laid across genotypes (p = 0.081). On average, females deposited 3.0 ± 0.5 egg batches per plant over the 18-day period.

**Fig 1.**
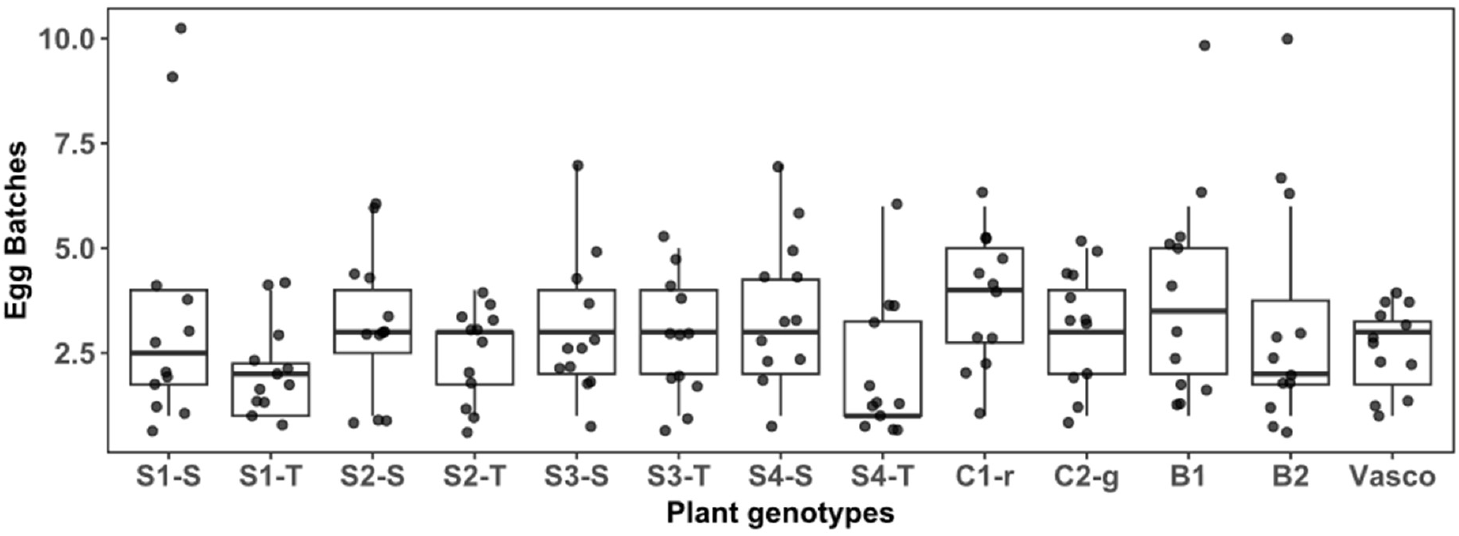
Oviposition preferences of *P. leporinus* females in a no-choice test. Groups of 10 newly emerged females and 5 males were exposed to a single *Beta vulgaris* genotype for 18 days, and the number of egg batches laid was recorded. Boxplots with overlaid individual data points show the number of egg batches on different genotypes: sugar beet (S1-S, S1-T, S2-S, S2-T, S3-S, S3-T, S4-S, S4-T), Swiss chard (red, C1-r; green, C2-g), beetroot (B1, B2), and the sugar beet cultivar used for laboratory rearing (Vasco). In each boxplot, the central line represents the median; boxes span the 25th–75th percentiles (interquartile range, IQR); and whiskers extend to the most extreme values within 1.5 × IQR. All individual data points are plotted as jittered dots to show the full distribution. Differences among genotypes were not significant (n.s., p > 0.05; n = 12 per genotype).

These results indicate that, in the absence of alternative host options, newly emerged *P. leporinus* adults oviposit readily and to a comparable extent on all evaluated *B. vulgaris* genotypes, including Swiss chard and beetroot.

### Oviposition choice test

In the three-choice test, *P. leporinus* females laid eggs on all tested *B. vulgaris* genotypes, but significant differences were observed in some genotypes (Fig 2 a–f).

**Fig 2.**
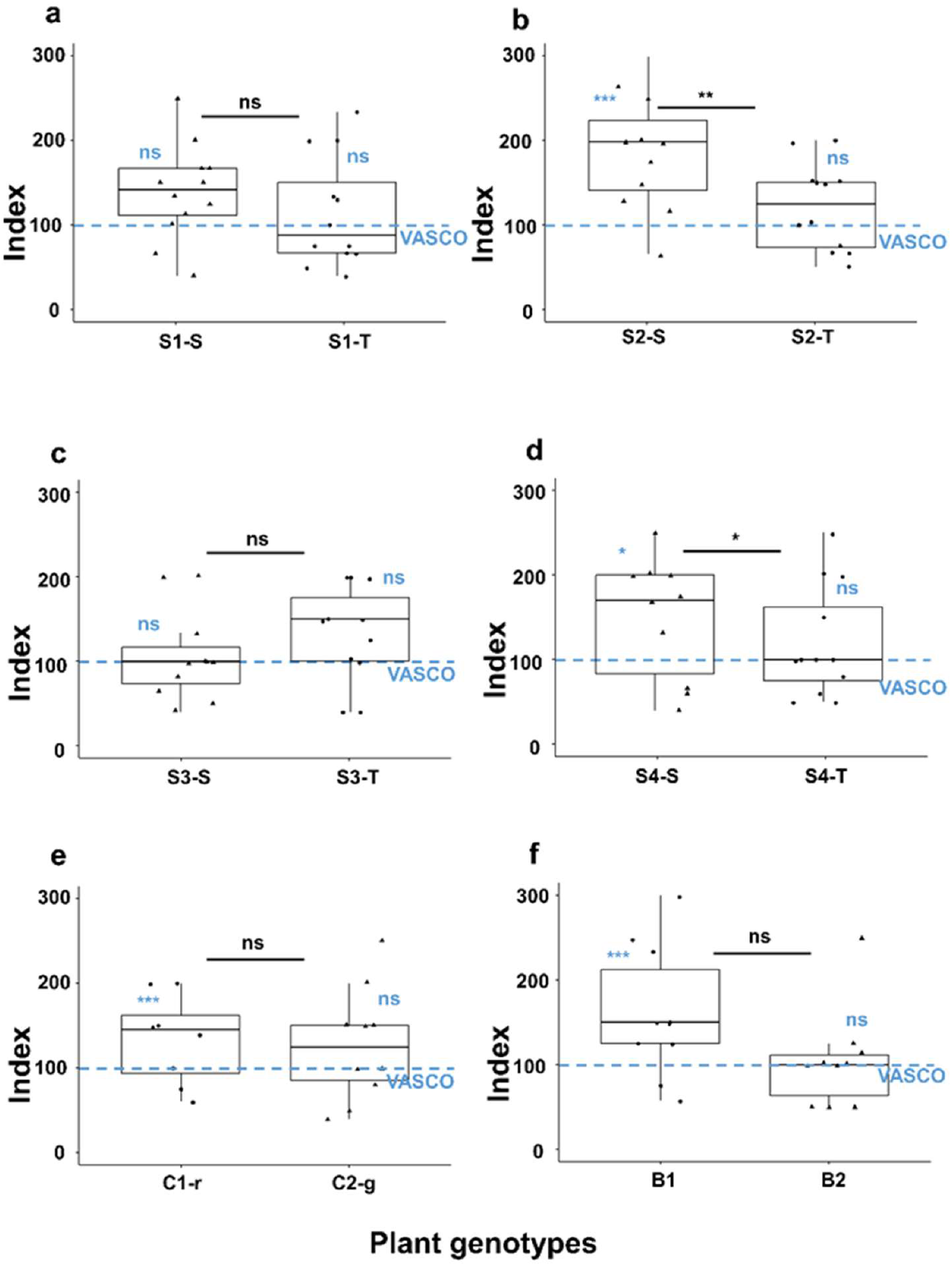
Oviposition preferences of *P. leporinus* in a three-choice test. Boxplots show the number of egg batches laid on different *B. vulgaris* genotypes by 10 newly-emerged females when exposed to the three-choice setup with 5 males for 18 days, after which egg batches per genotype were counted. Panels (a-f) correspond to comparisons of preference between two *B. vulgaris* genotypes (S1-S, S1-T, S2-S, S2-T, S3-S, S3-T, S4-S, S4-T), Swiss chard (red, C1-r; green, C2-g), beetroot (B1, B2), and the sugar beet variety used for the rearing (Vasco) as a control, which is shown as a baseline. The y-axis represents the choice index (0 = no choice). In the boxplots, the line inside each box shows the median, the box spans the 25th to 75th percentiles (interquartile range, IQR), and the whiskers extend to the smallest and largest values within 1.5 × IQR. Individual points beyond this range are also plotted as jittered dots to show all observations. Statistical comparisons are shown in black for differences between genotypes, and in blue for differences between each genotype and Vasco. Asterisks indicate significant differences: ***p < 0.001; **p < 0.01; *p < 0.05; not significant (n.s., p ≥ 0.05)

Females equally preferred genotypes S1-S, S1-T, and Vasco, as well as genotypes S3-S, S3-T, and Vasco, with no significant differences in oviposition (p > 0.05; Fig 2a, c).

In contrast, females deposited more egg batches on genotype S2-S (3.58 ± 0.52) than on genotype S2-T (1.33 ± 0.39) and Vasco (p = 0.0003 and p = 0.002, respectively), whereas oviposition on genotype S2-T did not differ from Vasco (p = 0.73; Fig 2b). Genotype S4-S (2.75 ± 0.50) received more egg batches than genotype S4-T (1.3 ± 0.4) and Vasco (p = 0.03 and p = 0.01, respectively), while genotype S4-T did not differ from Vasco (p = 0.98; Fig 2d).

Oviposition on Swiss chard genotypes C1-r and C2-g did not differ significantly from each other (p = 0.19). However, genotype C1-r (2.8 ± 0.6) received more eggs than Vasco (0.7 ± 0.4; p < 0.001), whereas genotype C2-g did not differ from Vasco (p = 0.052; Fig 2e).

Similarly, oviposition on beetroot genotypes B1 and B2 did not differ significantly (p = 0.066). Yet, genotype B1 (3.8 ± 0.8) attracted more egg batches than Vasco (1.3 ± 0.4; p < 0.001), while genotype did not differ from Vasco (p = 0.206; Fig 2f).

Taken together, these results indicate that *P. leporinus* females discriminate among *B. vulgaris* genotypes when choosing oviposition sites. Specific genotypes—particularly S2-S, S4-S, C1-r, and B1—were more attractive for egg laying, suggesting that genotype-specific plant traits influence adult host selection and oviposition behavior.

### Development of larvae on *Beta vulgaris* genotypes

*P. leporinus* nymphs exhibited comparable performance across different *B. vulgaris* genotypes (Fig 3). Final individual weight (Fig 3a) did not differ significantly among genotypes, indicating similar growth rates, with a final mean weight of 5.1 ± 0.1 mg per nymph. Similarly, over 85% of all individuals reached the fifth instar after 90 days of feeding (Fig 3b), with no significant differences observed among sugar beet (S1-S, S1-T, S2-S, S2-T, S3-S, S3-T, S4-S, S4-T), Swiss chard (red, C1-r; green, C2-g) and beetroot (B1, B2). Mortality rates (Fig 3c) were also consistent across all genotypes, with an overall average of 19.7 ± 1.1% after 90 days.

**Fig 3.**
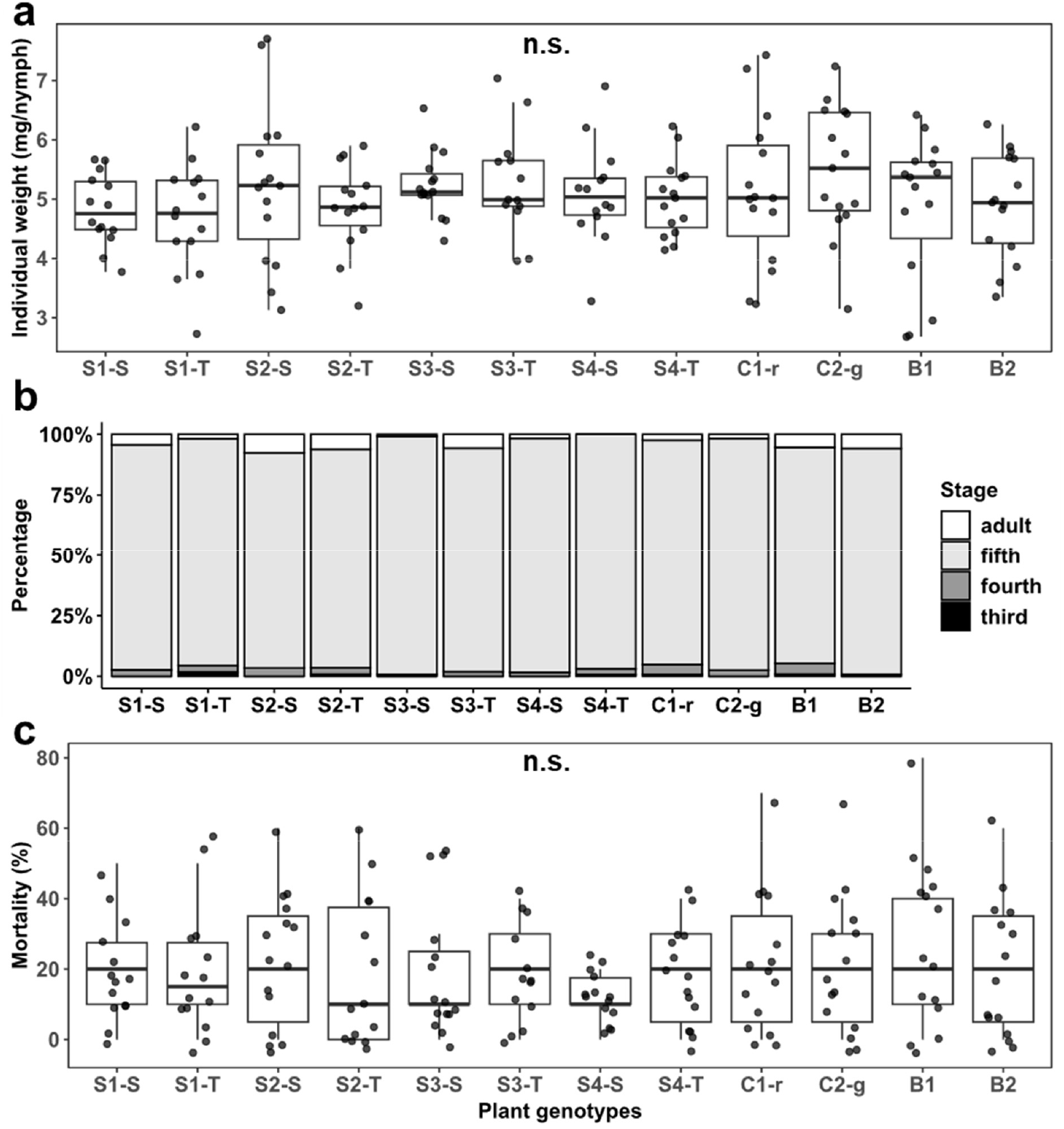
Performance of *P. leporinus* nymphs on different *Beta vulgaris* genotypes. (a) Final individual weight ± SE, (b) instar reached, and (c) mortality were measured for groups of ten second-instar nymphs after 90 days of feeding on a single plant of sugar beet (S1-S, S1-T, S2-S, S2-T, S3-S, S3-T, S4-S, S4-T), Swiss chard (red, C1-r; green, C2-g), or beetroot (B1, B2) (n = 12 per genotype).Boxplots show the median (horizontal line), interquartile range (box; 25th–75th percentiles), and whiskers extending to the most extreme values within 1.5 × IQR; all individual observations are plotted as jittered dots to display the full distribution. P values are given for treatments, obtained from generalized linear models with Gaussian distribution (final weight, instar reached) or binomial distribution (mortality). Differences among genotypes were not significant (n.s., p > 0.05).

Taken together, these results suggest that *P. leporinus* nymph development and survival are not significantly influenced by the *B. vulgaris* genotype on which they feed.

### Belowground olfactometer tests

Olfactometer assays showed that *P. leporinus* nymphs generally responded similarly to root volatiles of most *B. vulgaris* genotypes, although some significant differences were detected (Fig 4 a–f).

**Fig 4.**
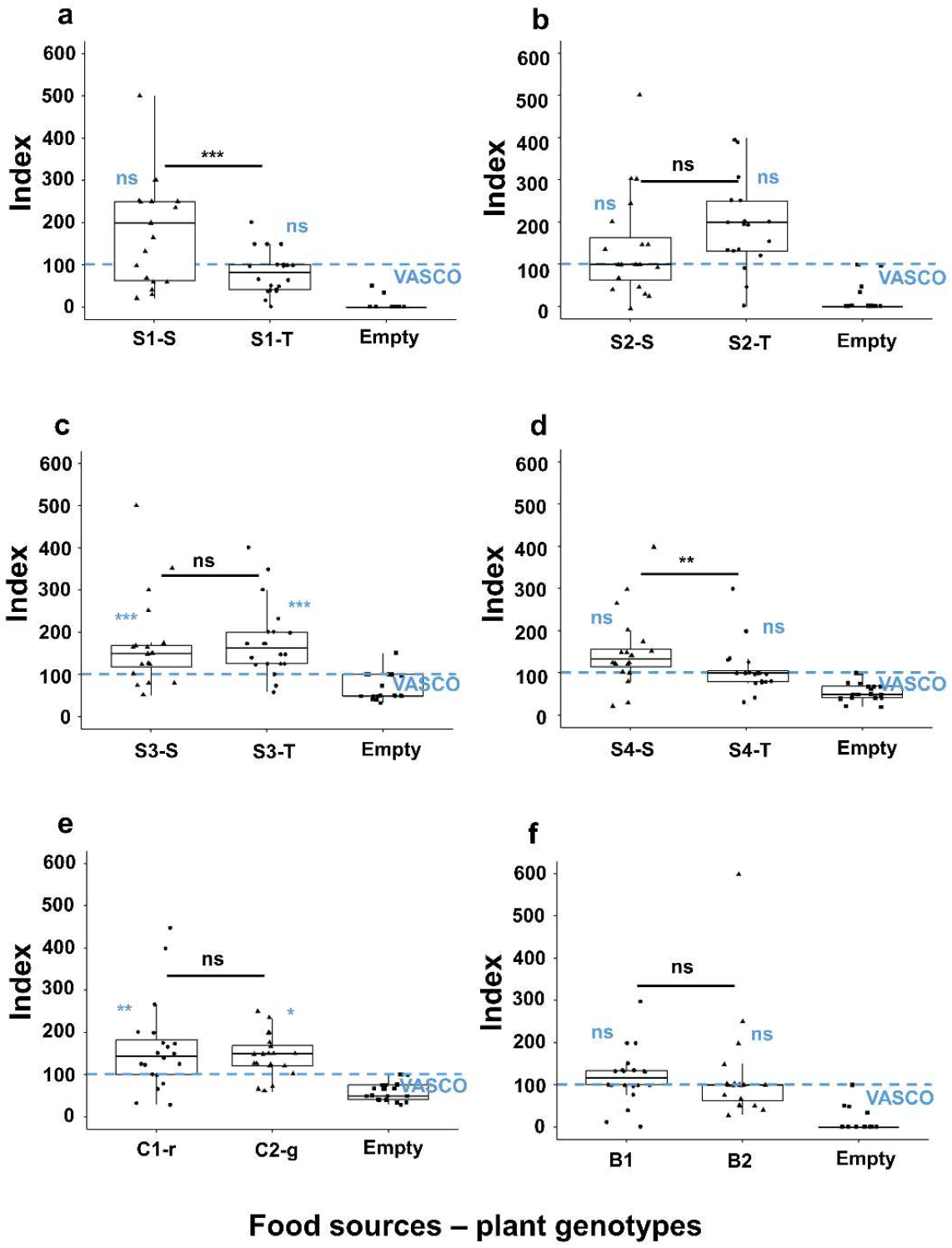
Olfactometer responses of *P. leporinus* nymphs to root volatiles from different *B. vulgaris* genotypes. Nymph olfactory preferences were assessed using a four-arm belowground olfactometer. Third-instar nymphs were introduced into the central chamber, and their positions were recorded after 24 hours. Panels (a-f) correspond to comparisons of preference between two *B. vulgaris* genotypes (S1-S, S1-T, S2-S, S2-T, S3-S, S3-T, S4-S, S4-T), Swiss chard (red, C1-r; green, C2-g), beetroot (B1, B2), an empty arm and the sugar beet variety used for the rearing (Vasco) as a control, which is shown as a baseline. The y-axis represents the choice index (0 = no choice). In the boxplots, the line inside each box shows the median, the box spans the 25th to 75th percentiles (interquartile range, IQR), and the whiskers extend to the smallest and largest values within 1.5 × IQR. Individual points beyond this range are also plotted as jittered dots to show all observations. Statistical comparisons are shown in black for differences between genotypes and in blue for differences between each genotype and Vasco, as well as the empty arm vs. Vasco. Asterisks are indicated in black between each pair of genotypes evaluated, and in blue for pairwise comparisons between each genotype and the baseline variety Vasco. Asterisks indicate significant differences: ***p < 0.001; **p < 0.01; *p < 0.05; not significant (n.s., p ≥ 0.05).

More nymphs were attracted to genotype S1-S (4.3 ± 0.4) compared to genotype S1-T (2.0 ± 0.2), while neither differed significantly from Vasco (p = 0.056 and p = 0.062, respectively; Fig 4a).

No differences were detected between genotypes S2-S and S2-T, or when compared with Vasco (p > 0.05; Fig 4b). A similar pattern was observed for beetroot genotypes B1 and B2, and also when compared with Vasco (p > 0.05; Fig 4f).

Genotypes S3-S (3.3 ± 0.3) and S3-T (3.9 ± 0.3) did not differ from each other, but both attracted more nymphs than Vasco (1.7 ± 0.2; p = 0.001 and p < 0.001, respectively; Fig 4c).

Genotype S4-S (3.8 ± 0.4) attracted more nymphs than genotype S4-T (2.3 ± 0.3), whereas neither genotype differed significantly from Vasco (p = 0.840 and p = 0.087, respectively; Fig 4d).

Swiss chard genotypes C1-r (3.6 ± 0.4) and C2-g (3.4 ± 0.2) did not differ from each other, but both attracted significantly more nymphs than Vasco (2.1 ± 0.3; p = 0.007 and p = 0.020, respectively; Fig 4e).

Across all assays, Vasco was consistently preferred over the empty control arms, with the preference for Vasco.

These findings indicate that while most *B. vulgaris* genotypes elicited similar olfactory responses, specific genotypes, particularly S1-S, S3-S, S3-T, S4-S, C1-r, and C2-g, significantly influenced host location behavior in *P. leporinus* nymphs.

## Discussion

This study examined the interactions between *Pentastiridius leporinus* and twelve *Beta vulgaris* genotypes, focusing on oviposition behavior, nymphal development, and belowground chemotaxis. Adult females laid similar numbers of egg batches across genotypes in no-choice setups, suggesting broadly suitable conditions for oviposition. However, in choice experiments, females preferred certain genotypes, indicating more subtle variation in host attractiveness. Despite these behavioral differences, nymphal performance did not vary significantly across genotypes in terms of mortality, final weight, or developmental stage over 90 days. In belowground olfactometer assays, however, nymphs displayed marked preferences for the root volatiles of specific genotypes, suggesting that chemical cues influence host location during development. These preferences may reflect genotype-specific differences in volatile emissions, which merit further investigation to clarify their role in host selection.

Despite the lack of performance differences, certain genotypes may be related to fitness-relevant traits that manifest in later life stages or in field conditions. Importantly, the panel included both commercial cultivars and preselected lines with contrasting SBR susceptibility, indicating that nymphal performance alone may not directly correlate with disease tolerance traits (Clark & Messina, 1998; Gripenberg et al., 2010).

From a breeding perspective, additional performance parameters such as adult emergence, longevity, and reproductive success could reveal genotype-related effects not captured in our assays. These traits may be particularly relevant given the physiological constraints of *P. leporinus* as an herbivore that feeds on plants from different families. Generalists are often less capable of metabolizing plant-specific secondary compounds compared to specialists (Gols et al., 2008; Lampert, 2012; Poelman et al., 2008; Wittstock et al., 2004). Compounds such as betaines, betalains, and flavonoids, which are known to vary across *B. vulgaris* genotypes (Hanson & Wyse, 1982; Oh et al., 2022; Sokolova et al., 2024), could subtly influence vector fitness. For instance, the red Swiss chard (C1-r) and the two red beetroot genotypes (B1 and B2) likely contain elevated levels of these pigments, which may affect insect development or reproduction in ways not reflected by early-stage performance. Identifying such long-term or sublethal effects could inform selection of sugar beet lines that reduce vector fitness and SBR risk (Ali & Agrawal, 2012; Müller et al., 2001)

In belowground olfactometer assays, *P. leporinus* nymphs showed clear chemotactic preferences for specific *B. vulgaris* genotypes. To our knowledge, this is the first report of chemotactic behavior in the nymphal stage of this species. This ability to respond to root-emitted volatiles may play a role in host location within the rhizosphere, particularly after sugar beet harvest when nymphs must seek alternative hosts (Bressan et al., 2011). Similar mechanisms have been described in other systems, such as the western corn rootworm (*Diabrotica virgifera virgifera*), which is attracted to maize genotypes emitting (*E*)-β-caryophyllene (Robert et al., 2013). Although general resource-based cues, such as CO_2_ emissions associated with greater root biomass, can influence belowground insect behavior (Arce et al., 2021; Johnson & Gregory, 2006; Johnson & Nielsen, 2012), this factor was controlled in our design by using plants of comparable size and developmental stage. Thus, the observed preferences likely reflect differences in volatile profiles among the evaluated genotypes. Volatile-guided host location is well-documented in coleopteran root feeders, but it remains largely unstudied in hemipteran root herbivores (Johnson & Gregory, 2006; Johnson & Nielsen, 2012).

Although our olfactometer experiments used intact plants, volatile emissions can change dramatically upon herbivore damage (Holopainen & Blande, 2013; War et al., 2011). Exploring how herbivore-induced root volatiles vary among *B. vulgaris* genotypes could clarify whether infested roots emit cues that attract conspecific nymphs. For instance, during the performance assays we observed gregarious behavior, with nymphs clustering in wax-coated nests built from their abdominal secretions. These waxes may serve antimicrobial or defensive functions (Lucchi & Mazzoni, 2004), and aggregation may provide survival benefits. If nymphs are more attracted to roots already colonized by conspecifics, volatiles emitted from infested roots might serve as cues to locate and maintain social aggregations, which could confer fitness benefits (Robert et al., 2012).

In parallel, plant infection status may also alter volatile signals in ways that affect vector behavior. Prior studies have shown that SBR-infected sugar beet plants emit altered volatile profiles, as increased limonene, nonadecane, and an unidentified compound, and reduced toluene (Kais et al., 2023). These changes may attract *P. leporinus*, facilitating bacterial dispersal, as has been shown in other plant-pathogen-vector systems (Franco et al., 2021; Keesey et al., 2017; MacLean et al., 2014; Mauck et al., 2012; Mauck et al., 2010). Investigating how both herbivory- and infection-induced shifts in plant chemistry shape vector behavior could help identify chemical traits that breeders can select for when developing sugar beet cultivars less attractive or suitable to *P. leporinus*, thereby supporting more durable SBR resistance (Karban, 2011; War et al., 2012).

While the nymphs used in our performance and olfactometer experiments were not the direct offspring of ovipositing females, our data suggest that adult female preference for certain genotypes did not necessarily correspond to improved nymphal performance. Similarly, nymphal chemotactic responses did not consistently align with adult oviposition choices. Such mismatches between preference and performance are well documented in generalist herbivores (Gripenberg et al., 2010; Mayhew, 2001; Scheirs et al., 2003) and underscore the complexity of host selection. This has important implications for breeding strategies, as plants that are unattractive to adult *P. leporinus* may still support nymphal development. Indeed, Pfitzer et al. (2022) demonstrated that female adults did not lay eggs when exposed to wheat plants, despite wheat roots providing a suitable food source for nymphal development.

A better understanding of these behavioral and physiological interactions is essential for developing sugar beet genotypes that are less attractive or suitable for *P. leporinus*. Identifying traits that reduce vector performance, either through direct effects on development or via disrupted host location, can support integrated pest and disease management strategies. Future work should focus on genotype-specific differences in both constitutive and induced plant metabolites, especially in the context of herbivory and infection.

## Data availability

The data supporting the findings of this study are available from the data repository GRO.data: https://doi.org/10.25625/2VEPGG

## Author contributions

Conceptualization: PB, SR, KB, BK, JG, MR; Methodology: PB, SR, MR; Investigation: PB, SR; Formal analysis: PB, SR; Supervision: MR; Funding acquisition: MR, JG; Visualization: PB, SR; Project administration: MR; Writing - original draft: PB; Writing – reviewing & editing: SR, MR, KB, BK, JG

## Acknowledgements

We thank Katrin Waldstein and Jutta Schaper for rearing plants and insects. Technical support was also provided by Ruth Pilot, Jonas Watterott, Fruzsina Ficze and Marieke Bode. Furthermore, we thank Mirko Rakoski (Gemeinschaft zur Förderung von Pflanzeninnovation e. V.), Kerstin Krüger and Mario Schumann (KWS KGaA), Maria Köhler (Strube D&S GmbH), Niels Wynant and Heinrich Reineke (SESVanderHave), as well as Britt-Louise Lennefors (United Beet Seeds) for providing seed material and fruitful discussions.

